# Selection and subsequent physiological characterisation of industrial *Saccharomyces cerevisiae* strains during continuous growth at sub- and- supra optimal temperatures

**DOI:** 10.1101/2020.01.30.926709

**Authors:** Ka Ying Florence Lip, Estéfani García-Ríos, Carlos E. Costa, José Manuel Guillamón, Lucília Domingues, José Teixeira, Walter M. van Gulik

## Abstract

A phenotypic screening of 12 industrial yeast strains and the well-studied laboratory strain CEN.PK113-7D at cultivation temperatures between 12 °C and 40 °C revealed significant differences in maximum growth rates and temperature tolerance. Two *Saccharomyces cerevisiae* strains, one performing best at sub-, and the other at supra-optimal temperatures, plus the laboratory strain, were selected for further physiological characterization in well-controlled bioreactors. The strains were grown in anaerobic chemostats, at a fixed specific growth rate of 0.03 h^-1^ and sequential batch cultures at 12, 30, and 39 °C. We observed significant differences in biomass and ethanol yields on glucose, biomass protein and storage carbohydrate contents, and biomass yields on ATP between strains and cultivation temperatures. Increased temperature tolerance coincided with higher energetic efficiency of cell growth, indicating that temperature intolerance is a result of energy wasting processes, such as increased turnover of cellular components (e.g. proteins) due to temperature induced damage.

## 1. Introduction

The alcoholic beverage and bio-ethanol industries mainly use *Saccharomyces* yeasts as their workhorses, because of their robustness to low pH and high ethanol tolerance. The ethanol yield and productivity of fermentation processes highly depend on the performance of the yeast strains used at the temperatures applied in these processes. Large differences in performance and adaptation to working temperatures exist between individual yeast strains [1].

Temperature is one of the predominant factors determining the operational costs of industrial fermentation processes. According to an energy study of the European Commission, the alcoholic beverage and bio-energy industries spend around 30-60 % of their total energy requirement of the whole production process to control the cultivation temperature [2]. In general, the optimum growth temperature of *Saccharomyces* yeasts lie between 28 °C to 33 °C [3]. However, this temperature range is not applicable for both the alcoholic beverage and bio-ethanol production processes in industry. In particular, wine fermentation processes are commonly operated at sub optimal temperatures (10-15 °C), mainly in white and rosé wines, to enhance and to retain their flavour volatiles [4]. These low working temperatures result in prolonged fermentation process duration and cause high risk of halted or sluggish fermentation [5]. Conversely, biofuel production processes are preferably performed at temperatures ≥40 °C especially for fermentation processes with simultaneous saccharification of lignocellulosic feedstocks [6, 7]. Therefore, the adaptation of yeast strains to temperatures outside the optimum range for growth provides an opportunity to make the production process more economical and eco-efficient.

Temperature tolerance is a polygenic trait which is influenced by a group of non-epistatic genes [8, 9]. Several studies have been performed to increase the understanding of the impact of the cultivation temperature on the physiology of *Saccharomyces* yeasts and to elucidate the mechanisms which contribute to differences in temperature tolerance [10–22]. In the majority of these studies, temperature shocks were applied rather than prolonged temperature stress, while the latter is much more relevant for industrial processes. To understand the cellular response and adaptation to temperature, the chosen research methodology is crucial to dissect transient stress responses and adaptation. This work aims at addressing the long term impacts of different cultivation temperatures on *Saccharomyces* strains with better growth performance at sub- and-supra optimal temperatures. We first characterized a collection of industrial *Saccharomyces* strains in terms of their capabilities to grow at sub-and-supra optimal temperatures ranging from 12 °C to 40 °C. This allowed us to select one strain which performed best at sub-optimal and another strain which performed best at supra-optimal temperatures. Subsequently, the physiological responses of these strains, together with a well-characterized laboratory strain, CEN.PK113-7D, to sub-optimal, optimal and supra-optimal temperatures were investigated in well-defined chemostat cultures at a constant growth rate.

## 2. Results

### 2.1. Growth phenotypic comparison of industrial Saccharomyces strains at different temperatures (12 °C to 40 °C)

The growth capacities of 13 *Saccharomyces* yeasts were determined at temperatures between 12 °C and 40 °C in aerobic microtiter plate cultivations. For all cultivations the purity of the strains was verified through analysis of DNA delta sequences after PCR amplification as described previously [23] (data not shown). Via this growth phenotypic screening, an inventory was made of the tolerance of these strains to the sub-optimal temperatures used in the alcoholic beverage industry and the supra-optimal temperatures used in the bio-fuel production industry. Due to the Crabtree effect [24], all strains showed diauxic growth in the presence of oxygen. Only the initial parts of the growth curves, representing growth on glucose with concurrent production of ethanol, were used. The obtained growth profiles of the strains at the different temperatures were fitted to the corrected modified Gompertz model as proposed by Zwietering *et al*. [25] which was modified from Salvado *et al*. [3]. The fit of this model to the experimental data yielded two parameters, a maximum specific growth rate (µ_max_) and a lag time (λ). The obtained lag times of the strains at the different cultivation temperatures were not correlated with the µ_max_ (data not shown). The fit of the model to the data was satisfactory for all strains at all cultivation temperatures with R-squared values (R^2^) ranging from 0.92 to 0.99.

A hierarchical cluster analysis (HCL) of the relation between µ_max_ and cultivation temperature was performed using Euclidean distance to further analyse the growth performance between different strains within the temperature range from 12°C to 40°C (Figure 1). The colours in the heat map represent the values of µ_max_. Based on the growth performance of all strains, the HCL dissected the applied temperature ranges into a range with sluggish growth (blue) and with facilitated growth (black or yellow). At 12 °C, 15 °C, and 40 °C, the growth rates of all strains were below the median (0.25 h^-1^). The cluster wherein facilitated growth occurred could be divided into two subgroups (optimal and sub- or-supra optimal growth). The optimal growth temperatures of all strains were observed at 28 °C and 33 °C, at which the heat map at this area mostly shows a yellow colour. The sub- and-supra optimal growth occurred at 25 °C and below and 37 °C and above, respectively, at which the heat map mostly shows a black colour. Regarding the sluggish and facilitated growth conditions, the HCL divided all strains into two major groups (1 and 2). Strains in group 1 had poor growth performance at all growth conditions, whereas strains in group 2 had a comparatively better growth performance.

**Figure 1.**
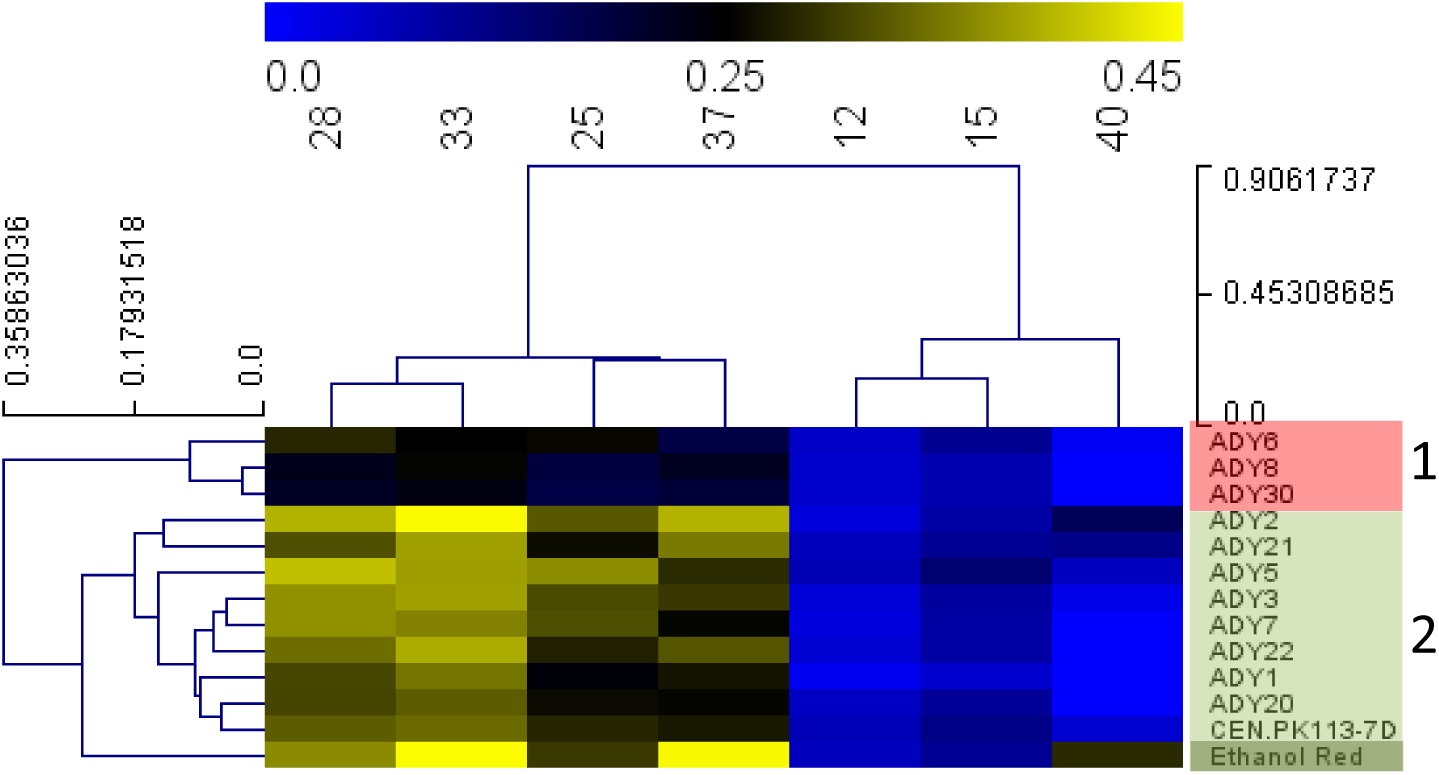
Heat map representing the HCL analysis using Euclidean distance with the area under the curve (AUC) of the 13 Saccharomyces yeasts. The data was obtained from the phenotypic screening experiment in microtiter plates. Growth rate range with sluggish growth is represented in blue, while facilitated growth is represented in black or yellow. Strains categorized in red (group 1) had slower growth rate at all growth conditions, while those in green (group 2) had faster rate at all growth conditions.

To obtain relations for the µ_max_ as a function of the cultivation temperature for all strains tested, the growth rates obtained for the different strains at the different cultivation temperatures were used to fit the cardinal temperature model with inflection point (CTMI) [3]. The goodness of fit of the CTMI model to the µ_max_ data was satisfactory in all cases, with p-values between 0.97 and 0.99. The CTMI fits of µ_max_ vs growth temperature for the different strains are shown in Figure 2. ADY5 clearly showed the fastest growth in the temperature range from 12 °C to 27 °C. At temperatures between 27 °C and 33 °C, ADY2 had the highest µ_max_, while between 33 °C and 40 °C Ethanol Red grew faster than all other strains. Ethanol Red and ADY5 respectively showed the best thermo- and cryo-tolerance, respectively, and were selected for further investigation of the underlying molecular and metabolic mechanisms. ADY5 is a *S. cerevisiae* x *S. cerevisiae* hybrid strain which is particularly used for the production of aromatic white and rosé wines in the industry at low fermentation temperatures and low nitrogen levels. Ethanol Red is an industrial yeast strain with high ethanol tolerance and is commonly used for the production of industrial ethanol at fermentation temperatures up to 40 °C.

**Figure 2.**
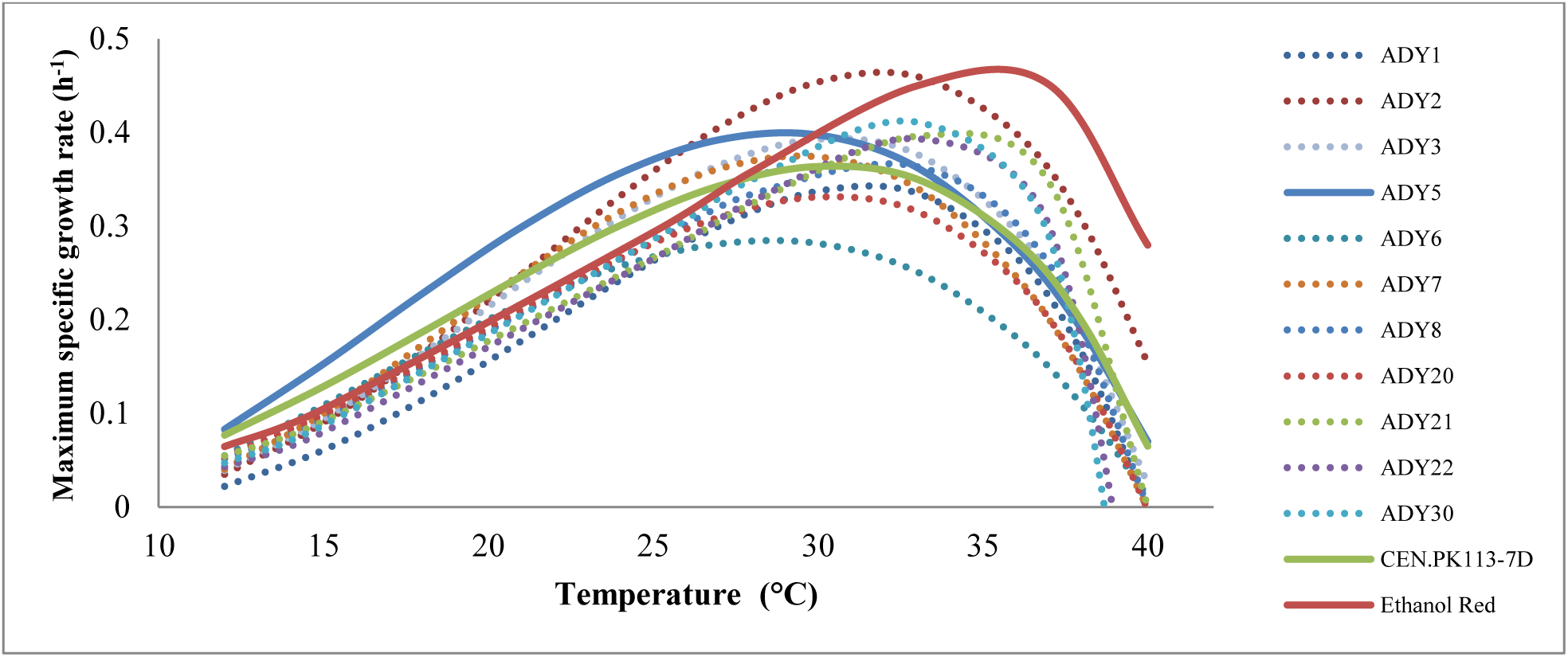
CTMI fit of the maximum specific growth rate as a function of the cultivation temperature (from 12 °C to 40 °C) to all of the 13 *Saccharomyces* yeasts.

The CTMI fit also provided the cardinal growth parameters of each strain (T_max_, T_opt_, T_min_, and μ_opt_). The average values of the estimated parameters for all strains were obtained from three independent experiments and are summarized in Table 1. The T_min_ of all strains ranged from 1 °C to 8 °C, whereas the T_max_ of all strains ranged from 39 °C to 41 °C. For all 13 strains, the optimum growth temperatures, T_opt_, were in the range between 29 °C and 35 °C whereby the corresponding specific growth rates ranged from 0.30 h^-1^ to 0.46 h^-1^.

**Table 1.** Estimated parameters with standard deviation from the CTMI fit for all the 13 strains. Standard deviations for each parameter were obtained from three independent non-linear fits.

| Strains | $\mu_{\text{opt}} (\text{h}^{-1})$ | $T_{\text{max}} (^{\circ}\text{C})$ | $T_{\text{min}} (^{\circ}\text{C})$ | $T_{\text{opt}} (^{\circ}\text{C})$ |
| --- | --- | --- | --- | --- |
| ADY1 | $0.343 \pm 0.008$ | $39.97 \pm 0.01$ | $7.78 \pm 0.45$ | $31.45 \pm 0.02$ |
| ADY2 | $0.464 \pm 0.011$ | $41.58 \pm 0.56$ | $7.36 \pm 0.79$ | $31.86 \pm 0.25$ |
| ADY3 | $0.381 \pm 0.009$ | $41.02 \pm 0.41$ | $7.29 \pm 0.09$ | $29.61 \pm 0.39$ |
| ADY5 | $0.409 \pm 0.002$ | $40.61 \pm 0.02$ | $4.06 \pm 0.39$ | $29.17 \pm 0.65$ |
| ADY6 | $0.297 \pm 0.002$ | $40.12 \pm 0.02$ | $4.14 \pm 0.00$ | $29.38 \pm 0.09$ |
| ADY7 | $0.375 \pm 0.004$ | $39.98 \pm 0.01$ | $7.34 \pm 0.90$ | $29.47 \pm 0.51$ |
| ADY8 | $0.367 \pm 0.008$ | $40.01 \pm 0.00$ | $3.93 \pm 0.00$ | $31.94 \pm 0.59$ |
| ADY20 | $0.330 \pm 0.006$ | $39.97 \pm 0.01$ | $4.79 \pm 0.84$ | $30.43 \pm 0.12$ |
| ADY21 | $0.302 \pm 0.010$ | $40.00 \pm 0.00$ | $2.11 \pm 0.75$ | $35.00 \pm 0.17$ |
| ADY22 | $0.370 \pm 0.027$ | $39.98 \pm 0.02$ | $1.79 \pm 0.93$ | $32.50 \pm 0.13$ |
| ADH30 | $0.236 \pm 0.004$ | $40.00 \pm 0.00$ | $4.00 \pm 0.10$ | $31.77 \pm 0.64$ |
| CEN.PK113-7D | $0.368 \pm 0.018$ | $41.21 \pm 0.81$ | $3.08 \pm 0.92$ | $30.03 \pm 0.85$ |
| Ethanol Red | $0.467 \pm 0.004$ | $41.48 \pm 0.05$ | $1.45 \pm 0.09$ | $35.26 \pm 0.18$ |

### 2.2. Physiological characterization of the selected strains at sub-optimal, optimal and supra-optimal temperatures

#### 2.2.1. Maximum specific growth rate determination of the selected strains in anaerobic sequential batches cultures

To validate the phenotypic screening results and the maximum growth rate estimations from the CTMI at the different temperatures for the selected strains (ADY5 and Ethanol Red) and the reference strain (CEN.PK113-7D), we performed sequential batch reactor (SBR) cultivations of the three strains. The carbon dioxide production profiles during the exponential phases of these SBR cultivations were used to calculate the µ_max_ of the selected strains at the three different cultivation temperatures (Table 2). Ethanol Red grew the fastest at 39 °C, whereas ADY5 grew the fastest at 12 °C, thereby confirming the results from the screening experiments in microtiter plates. Compared to these strains, CEN.PK113-7D showed the lowest maximum specific growth rate at all cultivation temperatures.

**Table 2.**
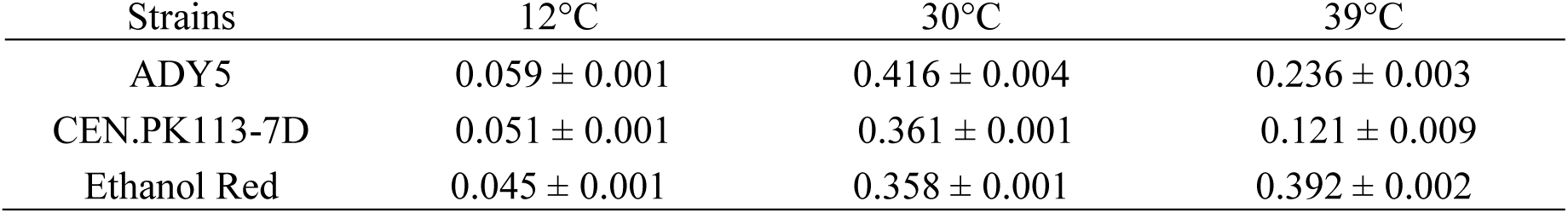
Maximum specific growth rates (h^-1^) in anaerobic SBR cultivations at 12 °C, 30 °C, and 39 °C. Standard errors for each strain and for each growth temperature were obtained from three independent batches.

| Strains | 12°C | 30°C | 39°C |
| --- | --- | --- | --- |
| ADY5 | $0.059 \pm 0.001$ | $0.416 \pm 0.004$ | $0.236 \pm 0.003$ |
| CEN.PK113-7D | $0.051 \pm 0.001$ | $0.361 \pm 0.001$ | $0.121 \pm 0.009$ |
| Ethanol Red | $0.045 \pm 0.001$ | $0.358 \pm 0.001$ | $0.392 \pm 0.002$ |

#### 2.2.2. Further physiological characterization of the strains in anaerobic chemostat cultures

To identify possible underlying mechanisms for the superior growth performances of ADY5 and Ethanol Red at respectively sub- and-supra optimal temperatures, we compared their physiology at 12 °C, 30 °C, and 39 °C with the well-studied laboratory strain CEN.PK113-7D under well-defined conditions at a constant specific growth rate. To this end the strains were grown at these three temperatures in anaerobic steady-state chemostat cultures at a dilution rate of 0.03 h^-1^. This dilution rate was slightly below the µ_max_ of the selected strains as well as CEN.PK113-7D at 12°C under anaerobic conditions (Table 2). During all steady-states the measured residual glucose concentration was below 0.50 mmol L^-1^, confirming glucose limited conditions. Chemostat instead of batch cultivation was chosen as this allowed to dissect the temperature effects from the effects of the specific growth rate. It is well known that differences in growth rate result in physiological changes in yeast, such as transcript levels [4]. Fully anaerobic instead of micro-aerobic conditions were chosen to rule out effects of differences in dissolved oxygen levels at different cultivation temperatures. Besides, alcoholic beverage and bio-fuel production is mainly performed in the absence of oxygen [26, 27].

#### 2.2.3. Large differences between strains in the effect of temperature on net conversion rates

The biomass specific conversion rates (q_i_, mmol·g_DW_ ·h) were calculated for all steady-state chemostat cultivations at the three different temperatures. The first order evaporation constants for culture broth and ethanol during chemostat cultivations at different temperatures were experimentally determined (Table S3, supplementary) and used for a proper calculation of the specific ethanol production rate. Simultaneous metabolic flux analysis and data reconciliation was applied (see materials and methods) and yielded the best estimations of the conversion rates within their error margins (Table 3 and Table 4). The reconciled data for the nine chemostat conditions fitted well with the experimental data as the p-values of the reconciled conversion rates were all greater than the significance level of 0.05 (data not shown).

**Table 3.** Reconciled specific conversion rates of the three strains with their standard errors during anaerobic chemostat cultivation at 12 °C, 30 °C, and 39 °C at a dilution rate of 0.03 h^-1^. The nomenclature of s, eth, and gly represents as substrate (glucose), ethanol, and glycerol, respectively.

| | $q_s$ (mmol·gDW<sup>-1</sup>·h<sup>-1</sup>) | $q_{CO_2}$(mmol·gDW<sup>-1</sup>·h<sup>-1</sup>) | $q_{eth}$ (mmol·gDW<sup>-1</sup>·h<sup>-1</sup>) | $q_{gly}$ (mmol·gDW<sup>-1</sup>·h<sup>-1</sup>) |
| --- | --- | --- | --- | --- |
| ADY5 |  |  |  |  |
| 12°C | -2.749 ± 0.100 | 4.560 ± 0.201 | 4.339 ± 0.201 | 0.428 ± 0.011 |
| 30°C | -2.078 ± 0.052 | 3.568 ± 0.104 | 3.399 ± 0.105 | 0.299 ± 0.011 |
| 39°C | -3.411 ± 0.007 | 4.805 ± 0.014 | 4.658 ± 0.014 | 0.309 ± 0.005 |
| CEN.PK113-7D |  |  |  |  |
| 12°C | -1.804 ± 0.092 | 3.055 ± 0.183 | 2.899 ± 0.183 | 0.298 ± 0.008 |
| 30°C | -1.761 ± 0.067 | 2.955 ± 0.134 | 2.794 ± 0.134 | 0.306 ± 0.008 |
| 39°C | -2.158 ± 0.049 | 3.558 ± 0.096 | 3.464 ± 0.097 | 0.270 ± 0.012 |
| Ethanol Red |  |  |  |  |
| 12°C | -1.594 ± 0.092 | 2.599 ± 0.184 | 2.418 ± 0.184 | 0.334 ± 0.006 |
| 30°C | -1.837 ± 0.048 | 3.030 ± 0.096 | 2.855 ± 0.096 | 0.326 ± 0.013 |
| 39°C | -2.069 ± 0.069 | 3.404 ± 0.138 | 3.242 ± 0.138 | 0.365 ± 0.009 |

**Table 4.** Reconciled specific conversion rates of the three strains with their standard errors during anaerobic chemostat cultivation at 12 °C, 30 °C, and 39 °C at a dilution rate of 0.03 h^-1^. The nomenclature of mal, suc, ace, and lac represents as malate, succinate, acetate, and lactate, respectively.

| | $q_{\text{mal}}$ (mmol·gDW<sup>-1</sup>·h<sup>-1</sup>) | $q_{\text{suc}}$ (mmol·gDW<sup>-1</sup>·h<sup>-1</sup>) | $q_{\text{ace}}$ (mmol·gDW<sup>-1</sup>·h<sup>-1</sup>) | $q_{\text{lac}}$ (mmol·gDW<sup>-1</sup>·h<sup>-1</sup>) |
| --- | --- | --- | --- | --- |
| ADY5 |  |  |  |  |
| 12°C | 0.008 ± 0.000 | 0.008 ± 0.000 | 0.067 ± 0.004 | 0.030 ± 0.004 |
| 30°C | 0.000 ± 0.000 | 0.000 ± 0.000 | 0.000 ± 0.000 | 0.080 ± 0.000 |
| 39°C | 0.007 ± 0.001 | 0.042 ± 0.003 | 0.016 ± 0.002 | 0.029 ± 0.002 |
| CEN.PK113-7D |  |  |  |  |
| 12°C | 0.006 ± 0.000 | 0.013 ± 0.000 | 0.000 ± 0.000 | 0.000 ± 0.000 |
| 30°C | 0.006 ± 0.006 | 0.013 ± 0.000 | 0.000 ± 0.000 | 0.000 ± 0.000 |
| 39°C | 0.014 ± 0.014 | 0.094 ± 0.000 | 0.021 ± 0.004 | 0.021 ± 0.004 |
| Ethanol Red |  |  |  |  |
| 12°C | 0.000 ± 0.000 | 0.013 ± 0.000 | 0.018 ± 0.001 | 0.018 ± 0.001 |
| 30°C | 0.000 ± 0.000 | 0.017 ± 0.000 | 0.012 ± 0.004 | 0.012 ± 0.004 |
| 39°C | 0.040 ± 0.040 | 0.001 ± 0.000 | 0.025 ± 0.001 | 0.025 ± 0.001 |

As expected from the stoichiometry of ethanol fermentation from glucose, the ratios (mol·mol^-1^) of the CO_2_ production rate (q_co2_) and ethanol production rate (q_eth_) were close to one for each strain and each temperature condition (Table 3). Although the dilution rate, and thus the specific growth rate, of all cultivations was the same, significant differences in the obtained q_i_ values were observed for different strains and at different cultivation temperatures. As an example, the specific glucose uptake rate (q_s_) of CEN.PK113-7D cultivated at the supra-optimal temperature (39 °C) was more than a factor of two higher than that of Ethanol Red cultivated at the sub-optimal temperature (12 °C). Similar differences were observed for the ethanol and carbon dioxide production rates. For all the strains, the glucose consumption as well as ethanol and CO_2_ production rates were highest at the highest cultivation temperature. However, the differences between the individual strains were large, whereby CEN.PK113-7D showed the highest values.

With respect to the sub-optimal temperature, the responses of the three strains were all different. For CEN.PK113-7D the q_s_, q_eth_, and q_CO2_ values were all significantly higher at 12 °C compared to the control temperature; ADY5 showed no significant differences while for Ethanol Red these specific conversion rates were all significantly lower at 12 °C compared to the control temperature (30 °C). These results clearly indicate large differences in cellular energetics between the distinct strains cultivated at different temperatures.

There were also differences in the production rates of glycerol (Table 3) and acidic by-products (Table 4) between the individual strains, but there appeared to be no clear correlation with the cultivation temperature. Glycerol production rates were similar for the different strains and temperatures, except for CEN.PK113-7D, which had a significantly increased glycerol production rate at the sub-optimal temperature, which was accompanied with an increased acetate production rate. Nevertheless, all three strains produced very small amounts of acids, with production rates up to 0.09 mmol·gdw ^-1^·h^-1^ (Table S1, supplementary).

#### 2.2.4. Large differences between strains in the effect of temperature on yields

A further analysis of the physiological differences between the strains grown at different temperatures was performed by comparing the yields of biomass and (by)products on glucose for the different cultivations (Table 5 and Table 6). Also, here, significant differences between strains and cultivation temperatures were observed. For all three strains the biomass yields on glucose were significantly lower at 39 °C compared to 30 °C, whereby CEN.PK113-7D had the lowest biomass yield. Likewise, at 12 °C CEN.PK113-7D had the lowest biomass yield. For ADY5 the biomass yields were the same at 12 °C and 30 °C, while for Ethanol Red the biomass yield was highest at 12 °C. The ethanol yields on glucose for the three strains at the different cultivation temperatures varied between 1.52 (Ethanol Red, 12 °C) and 1.70 (CEN.PK113-7D, 39 °C) mol ethanol per mol glucose, whereby each individual strain showed a slightly different ethanol yield on glucose in general (Table 5). Although Ethanol Red and ADY5 were designated, respectively, as hosts for the production of bioethanol and alcoholic beverages, they both had a lower ethanol yield on glucose than the laboratory strain CEN.PK113-7D, regardless of the cultivation temperature. CEN.PK113-7D showed the highest ethanol yield on glucose at 39 °C, which was accompanied with a corresponding low biomass yield on glucose. This clearly indicated that CEN.PK113-7D was negatively affected by the supra-optimal temperature which increased the cellular energy demand.

**Table 5.** Yields of biomass and main products on glucose of the three strains with their standard errors, calculated from the specific net conversion rates shown in Table 3.

| | $Y_{\text{biomass}}(\text{mol} \cdot \text{mol}^{-1})$ | $Y_{\text{CO}_2}(\text{mol} \cdot \text{mol}^{-1})$ | $Y_{\text{ethanol}}(\text{mol} \cdot \text{mol}^{-1})$ | $Y_{\text{glycerol}}(\text{mol} \cdot \text{mol}^{-1})$ |
| --- | --- | --- | --- | --- |
| ADY5 |  |  |  |  |
| 12°C | 0.595 ± 0.017 | 1.161 ± 0.046 | 1.607 ± 0.065 | 0.165 ± 0.005 |
| 30°C | 0.625 ± 0.014 | 1.422 ± 0.042 | 1.587 ± 0.049 | 0.174 ± 0.004 |
| 39°C | 0.514 ± 0.010 | 1.649 ± 0.096 | 1.605 ± 0.029 | 0.125 ± 0.003 |
| CEN.PK113-7D |  |  |  |  |
| 12°C | 0.396 ± 0.009 | 1.733 ± 0.050 | 1.649 ± 0.049 | 0.162 ± 0.005 |
| 30°C | 0.498 ± 0.011 | 1.717 ± 0.033 | 1.636 ± 0.032 | 0.144 ± 0.003 |
| 39°C | 0.404 ± 0.001 | 1.758 ± 0.003 | 1.704 ± 0.003 | 0.113 ± 0.001 |
| Ethanol Red |  |  |  |  |
| 12°C | 0.675 ± 0.020 | 1.630 ± 0.074 | 1.517 ± 0.072 | 0.210 ± 0.006 |
| 30°C | 0.601 ± 0.013 | 1.649 ± 0.034 | 1.554 ± 0.033 | 0.178 ± 0.004 |
| 39°C | 0.528 ± 0.011 | 1.645 ± 0.043 | 1.567 ± 0.042 | 0.176 ± 0.004 |

**Table 6.** Yields of by-products on glucose for the three strains with their standard errors, calculated from the specific net conversion rates shown in Table 4.

| | $Y_{\text{malate}}$ (mol·mol<sup>-1</sup>) | $Y_{\text{succinate}}$ (mol·mol<sup>-1</sup>) | $Y_{\text{acetate}}$ (mol·mol<sup>-1</sup>) | $Y_{\text{lactate}}$ (mol·mol<sup>-1</sup>) |
| --- | --- | --- | --- | --- |
| ADY5 |  |  |  |  |
| 12°C | 0.003 ± 0.000 | 0.007 ± 0.000 | 0.000 ± 0.000 | 0.000 ± 0.000 |
| 30°C | 0.004 ± 0.000 | 0.007 ± 0.000 | 0.000 ± 0.000 | 0.000 ± 0.000 |
| 39°C | 0.007 ± 0.000 | 0.044 ± 0.000 | 0.010 ± 0.001 | 0.021 ± 0.000 |
| CEN.PK113-7D |  |  |  |  |
| 12°C | 0.003 ± 0.000 | 0.003 ± 0.000 | 0.025 ± 0.001 | 0.011 ± 0.000 |
| 30°C | 0.000 ± 0.000 | 0.000 ± 0.000 | 0.000 ± 0.000 | 0.038 ± 0.001 |
| 39°C | 0.003 ± 0.000 | 0.016 ± 0.000 | 0.006 ± 0.000 | 0.011 ± 0.000 |
| Ethanol Red |  |  |  |  |
| 12°C | 0.000 ± 0.000 | 0.008 ± 0.000 | 0.011 ± 0.000 | 0.007 ± 0.000 |
| 30°C | 0.000 ± 0.000 | 0.009 ± 0.000 | 0.006 ± 0.001 | 0.033 ± 0.003 |
| 39°C | 0.019 ± 0.004 | 0.000 ± 0.000 | 0.012 ± 0.000 | 0.032 ± 0.001 |

#### 2.2.5. Temperature effect on metabolic efficiency of biomass formation

Metabolic flux analysis was performed for each individual strain at each cultivation temperature using a stoichiometric model for anaerobic growth of *S. cerevisiae* on glucose. Hereby the metabolic flux distributions were calculated using the biomass specific conversion rates obtained from the steady-state chemostat cultivations as input. This allowed to calculate the energetic efficiency of growth of each individual strain as a function of the cultivation temperature. For each condition, we calculated the net biomass specific rate of catabolic ATP production by summing up the hexokinase, glycerol-3-phosphase, phosphofructokinase, phosphoglycerate kinase, and pyruvate kinase fluxes. The ratios of the biomass specific growth rates and net biomass specific ATP production rates provided the biomass yields with respect to the produced ATP (Y_x/ATP_) at the different cultivation temperatures for each strain (Table 7). In spite of the fixed dilution rate, and thus identical specific growth rates, we observed large differences between the individual strains at the different cultivation temperatures. Ethanol Red cultivated at 12 °C produced three times more biomass per mole of ATP than CEN.PK113-7D cultivated at 39 °C. For each cultivation temperature, the Y_x/ATP_ values for Ethanol Red and ADY5 were significantly higher than for CEN.PK113-7D. Ethanol Red grew most efficiently at both 12 °C and 39 °C. A possible reason for the differences in growth efficiencies might be differences in biochemical composition, e.g. protein contents of the cells among distinct strains. Therefore, for all chemostat cultivations the cellular contents of protein and storage carbohydrates were quantified for each strain and cultivation temperature.

**Table 7.** Yields of biomass on ATP (g_DW_·mol_ATP_^-1^) with their standard errors for the anaerobic chemostat cultures.

|  | 12°C | 30°C | 39°C |
| --- | --- | --- | --- |
| ADY5 | $11.79 \pm 0.23$ | $12.75 \pm 0.18$ | $9.17 \pm 0.27$ |
| CEN.PK113-7D | $7.64 \pm 0.17$ | $9.45 \pm 0.12$ | $5.35 \pm 0.22$ |
| Ethanol Red | $15.24 \pm 0.30$ | $12.27 \pm 0.13$ | $10.69 \pm 0.16$ |

### 2.3. Cellular protein, glycogen and trehalose contents

During chemostat cultivation at 12 °C, the total cellular protein content was very similar for the three strains (Table 8), with an average value of about 0.32 g_protein_·g_DW_. At a cultivation temperature of 30 °C, the protein contents of ADY5 and Ethanol Red were slightly higher, while there was a significant increase for CEN.PK113-7D compared to the 12 °C cultivations. Remarkably, the total protein contents at the supra-optimal cultivation temperature of 39 °C were significantly lower for all strains. These results were confirmed by quantification of the total cellular nitrogen contents (Table S2, supplementary) which indeed showed the same trends.

**Table 8.** Total protein contents of biomass (g_protein_·g ^-1^) for the three strains with their standard deviations during anaerobic chemostat cultivation at different temperatures.

|  | 12°C | 30°C | 39°C |
| --- | --- | --- | --- |
| ADY5 | $0.316 \pm 0.004$ | $0.334 \pm 0.008$ | $0.274 \pm 0.030$ |
| CEN.PK113-7D | $0.332 \pm 0.010$ | $0.376 \pm 0.007$ | $0.246 \pm 0.002$ |
| Ethanol Red | $0.325 \pm 0.006$ | $0.342 \pm 0.005$ | $0.284 \pm 0.008$ |

**Table 9.** Yeast strains used in this study.

| Strains | Species | Origin |
| --- | --- | --- |
| ADY1 | *S.cerevisiae* | Lallemand Inc., France |
| ADY2 | *S.cerevisiae* | Lallemand Inc., France |
| ADY3 | *S.cerevisiae* | Lallemand Inc., France |
| ADY5 | *S.cerevisiae* x *S.cerevisiae* | Lallemand Inc., France |
| ADY6 | *S.uvarum* | Lallemand Inc., France |
| ADY7 | *S.cerevisiae* | Lallemand Inc., France |
| ADY8 | *S.cerevisiae* | Lallemand Inc., France |
| ADY20 | *S.cerevisiae* | Lallemand Inc., France |
| ADY21 | *S.cerevisiae* | Lallemand Inc., France |
| ADY22 | *S.cerevisiae* | Lallemand Inc., France |
| ADH30 | *S.cerevisiae* | Lallemand Inc., France |
| CEN.PK113-7D | *S.cerevisiae* | Fungal Biodiversity Centre, Utrecht, The Netherlands |
| Ethanol Red | *S.cerevisiae* | Fermentis, S.I. Lesaffre |

We observed larger differences for the cellular contents of glycogen and trehalose between different strains and cultivation temperatures (Figure 3 and Figure 4). For all cultivation temperatures, both Ethanol Red and ADY5 had higher glycogen and trehalose accumulations than CEN.PK113-7D. The accumulations of glycogen and trehalose for all three strains showed opposite patterns with respect to the cultivation temperature. All three strains had higher glycogen but lower trehalose accumulations at 12 °C, and vice versa at 39 °C. At 12 °C, the trehalose contents of all three strains were extremely low (below 1 %). We observed significant differences in glycogen accumulation between various strains at the sub-optimal temperature, whereby ADY5 had the highest value of about 20 %. At 39 °C, the trehalose accumulations of ADY5 and Ethanol Red were particularly high with values around 10 %. The glycogen accumulations at this temperature were also significantly different between strains. ADY5 had the highest value, more than 5 %, whereas CEN.PK113-7D had the lowest value, below 1 %.

**Figure 3.**
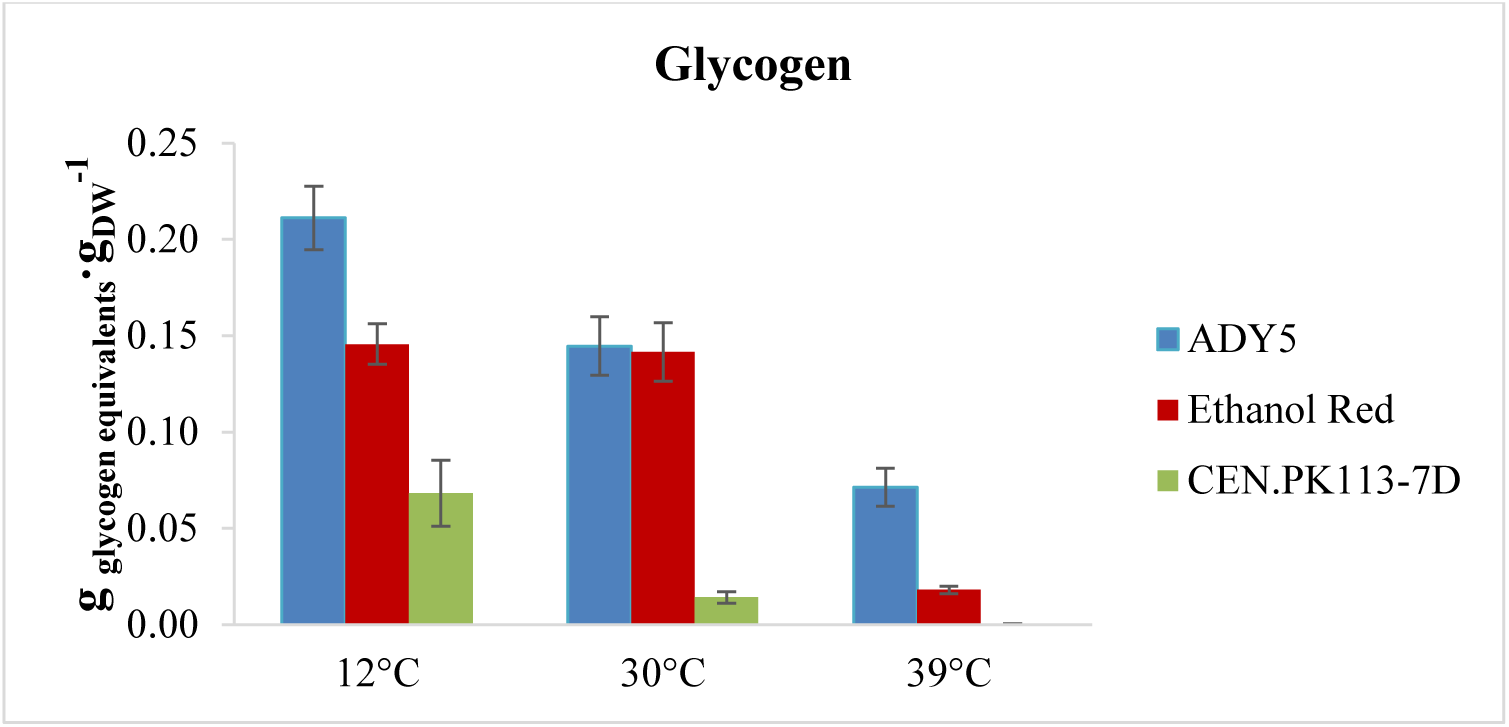
Glycogen accumulation of the three strains in anaerobic chemostat at 12 °C, 30 °C, and 39 °C. Error bars represent standard deviations of average values of measurements in biomass samples from identical chemostat cultures at four different time points in steady-state.

**Figure 4.**
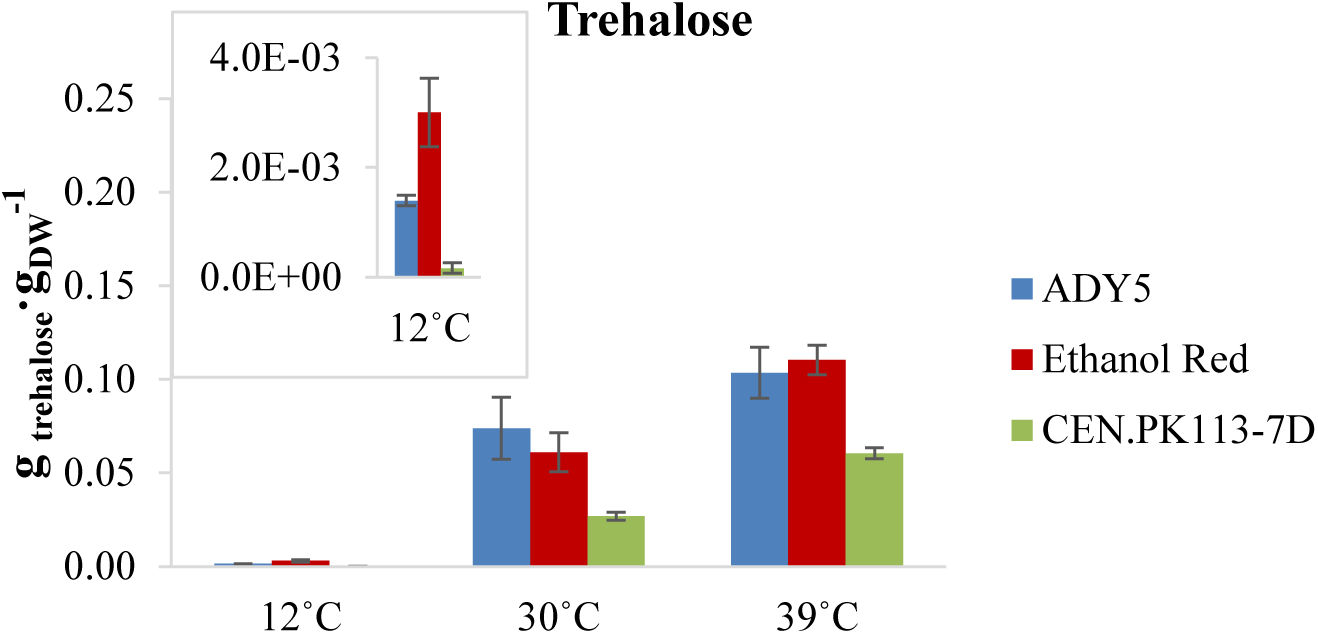
Trehalose accumulation of the three strains in anaerobic chemostat at 12 °C, 30 °C, and 39 °C. Error bars represent standard deviations of average values of measurements in biomass samples obtained from identical chemostat cultures collected at four different time points in steady-state.

## 3. Discussion

In this study we found that the impact of the cultivation temperature on the growth physiology of *Saccharomyces* yeasts is largely different between individual strains. A first growth phenotypic screening of one laboratory and 12 industrial yeast strains at growth temperatures ranging between 12 °C and 39 °C in microtiter plates, revealed significant differences in their optimum growth temperatures and temperature tolerance. From this initial screening of the strains, we selected the ones growing the fastest at sub- and supra-optimal temperatures (respectively ADY5 and Ethanol Red), plus the reference laboratory strain (CEN.PK113-7D), for in depth physiological characterization in anaerobic SBR and chemostat cultivations. We used SBR cultivation over normal batch cultivation because the SBR set-up was shown to give a better reproducibility and more consistent results [28] as the effect of carryover from the inoculum, which might play a role during the first batch, vanishes after a few repetitive batch cultivations. Within ten repetitive batches, we did not observe adaptation/evolution of cell cultures because no increase of the maximum specific growth rate occurred. The growth performance of these three strains at 12 °C, 30 °C and 39 °C in the anaerobic SBR cultures aligned well with the description of the CTMI model (Table 1) derived from the microtiter plate data. The μ_max_ values obtained from the anaerobic SBR cultivations (Table 2) showed that ADY5 and Ethanol Red clearly performed better at 12 °C and 39 °C, respectively, compared to CEN.PK113-7D. The μ_max_ values of ADY5 and CEN.PK113-7D at 30 °C were very similar to the estimated optimal μ_max_ obtained from the CTMI. The estimated optimum temperature of Ethanol Red from the CTMI was five degrees higher than that of the other two strains, highlighting its temperature tolerance, and was close to the value reported in the literature [29]. Therefore, the CTMI model was useful and reliable in our study to describe the growth profile of the selected strains over the temperature range.

Further physiological characterization of the three strains in anaerobic glucose limited chemostat cultures at a fixed dilution rate revealed very large differences in the biomass yields on glucose between the different strains, but also between the different cultivation temperatures for the same strain (Table 5). Generally, lower biomass yields correlated with higher ethanol and CO_2_ yields, suggesting differences in energy requirements for growth and maintenance for the different strains and temperatures. Also the formation of increased amounts of by-products (glycerol and acids) will result in decreased biomass yields. However, the total yield of by-products on glucose was very similar for all chemostat cultivations and was on average 0.100 ± 0.005 mol of carbon produced per mol of carbon consumed as glucose. Notably, CEN.PK113-7D, of which the biomass yields at the different temperatures were the lowest, also showed the lowest average by-product yields, indicating that by-product formation was not the cause of the low biomass yields. The observed large differences in biomass yields must therefore have been caused by large differences in cellular energy demands, which were quantified by calculating the biomass yields on ATP (Y_ATP_) (Table 7). The Y_ATP_ of anaerobically grown *S. cerevisiae* (CBS8066) has been determined previously from glucose limited chemostat experiments at a cultivation temperature of 30 °C [30] whereby the maximum value was 16 gDW·mol^-1^ ATP.

From retentostat experiments it was found that the ATP dissipation rate for maintenance (m_ATP_) of CEN.PK113-7D under anaerobic conditions equals 1 mmol_ATP_·gDW^-1^·h^-1^ at 30°C [31]. From these figures it can be calculated that Y_ATP_ should be around 14 gDW·mol^-1^ ATP at a growth rate of 0.1 h^-1^, which was indeed observed experimentally [30] and due to an increased contribution of maintenance energy requirements, around 10.5 gDW·mol^-1^ ATP at the growth rate of 0.03 h^-1^ used in our chemostat cultivations. The Y_ATP_ values we observed for the three strains at 30 °C (between 9.5 and 12.8 gDW·mol^-1^ ATP) are close to this value, whereby differences in biomass composition, especially protein content of which the biosynthesis is the most energy demanding, could be responsible for the differences in Y_ATP_ values between the strains at the same cultivation temperature. Quantification of the total protein contents revealed, however, that there were minor differences in protein contents between the three strains at the same temperature, thus ruling out that the observed differences in Y_ATP_ between the strains were caused by differences in protein content. The cultivation temperature itself had more effect, especially at 39 °C the protein contents of all three strains were significantly lower than at 30 °C and 12 °C. This could be a strategy of the cells to decrease their ATP expenses to cope with the stress during growth at supra-optimal temperatures.

It is well known that maintenance energy requirements of microorganisms increase with increasing cultivation temperature [32]. Using the correlation proposed by these authors, an increase of the cultivation temperature from 30 °C to 39 °C would result in a 2.2 fold increase of the maintenance coefficient, which would result in a decrease of Y_ATP_ with 30 % to 7.4 gDW·mol^-1^ ATP at a growth rate of 0.03 h^-1^. The decrease of Y_ATP_ of the ADY5 strain at 39 °C is indeed very close to 30% value while for CEN.PK113-7D the decrease is more than 40 %. Ethanol Red appeared clearly better adapted to higher cultivation temperatures as the Y_ATP_ decreased with only 13 %. Conversely, a decrease of the cultivation temperature from 30 °C to 12 °C would, according to the correlation of Tijhuis *et al*. [32], result in a decrease of the maintenance energy requirements with more than a factor of 5 and, consequently, an increase of Y_ATP_ with almost 40 % at a growth rate of 0.03 h^-1^. Such an increase was not observed in our chemostat cultures, on the contrary, for two strains (CEN.PK113-7D and ADY5) Y_ATP_ was lower at 12 °C than at 30 °C. Nevertheless for Ethanol Red Y_ATP_ was 24 % higher at 12 °C compared to 30 °C.

Another factor which can lead to differences in Y_ATP_ between different strains and/or cultivation temperatures is differences in the concentrations of weak acids [29]. Passive diffusion of the undissociated form into the cells and subsequent active export results in an ATP dissipating futile cycle leading to increased non-growth associated energy requirements. Of the acidic by products excreted, acetic acid (pKa = 4.76) would have the most significant influence on the maintenance energy requirements because at the cultivation pH of 5, 37 % of the acid is present in the undissociated form. The maximum residual acetic acid concentration of the chemostat cultures was 2.84 mmol·L^-1^ for CEN.PK113-7D cultivated at 12 °C (Table 12). At this residual acetic acid concentration, the maintenance energy requirements would be approximately 2.3 mmol_ATP_·gDW^-1^·h^-1^ at pH 5, and 30 °C [30] and would result in an Y_ATP_ of 7.2 gDW·mol^-1^ ATP, which is close to the observed value of 7.64 (Table 7). For all the other strains and temperatures the residual acetic acid concentrations were around 1 mM or lower and thus the effect on the maintenance energy requirements were assumed to be small. Interestingly, ADY5 did not produce acetic acid at 12 °C and 30 °C. Possible uncoupling of acetic acid seems not to attribute significantly to the Y_ATP_ of Ethanol Red at 12 °C where the residual acetic acid concentration was 2 times higher than that at 30°C.

Another cause of the effect of temperature on Y_ATP_ could be protein misfolding at high temperatures and aggregation at low temperatures. Remarkably the proteome analysis of the identical chemostat cultured strains revealed that for both CEN.PK113-7D and ADY5 the proteins related to protein folding and degradation processes were upregulated at 12 °C, in contrast to Ethanol Red [33]. This could indicate that protein aggregation and/or misfolding and subsequent degradation and re-synthesis might have occurred in these strains at 12 °C, resulting in an increased energy demand and thus a decreased Y_ATP_.

In Ethanol Red Erg13, one of the first and rate controlling enzymes in the ergosterol biosynthesis pathway was upregulated at both 12°C and 39°C. Although ergosterol was one of the anaerobic growth factors supplemented to the chemostat medium as its synthesis requires oxygen, this upregulation could indicate increased incorporation of ergosterol in the cell membrane of Ethanol Red. Several studies have reported that the activation of the ergosterol pathways makes yeast cells more resistant/tolerant to a variety of stresses, including low temperature, low-sugar conditions, oxidative stress and ethanol [34, 35].

All three strains showed upregulation of proteins involved in transport and metabolism of carbohydrates as well as energy and amino acid metabolism at 12 °C compared to 30 °C [33]. This shows that maintaining the same specific growth rate of 0.03 h^-1^ in the chemostat at 12°C, where maximum enzyme capacities have decreased, requires upregulation of proteins in central metabolism.

It is well-known that the accumulation of the storage carbohydrates glycogen and trehalose in *S. cerevisiae* strongly depends on the growth rate [36] and that in particular trehalose was shown to protect cells during stress conditions [37–39]. As in this work all cultivations were carried out at a fixed dilution rate, differences in storage carbohydrate accumulation can only be attributed to the particular strain used and/or the cultivation temperature. During glucose limited chemostat cultivation all three strains accumulated both trehalose and glycogen, whereby the differences in total accumulations (glycogen and trehalose) between strains were more significant than between cultivation temperatures for the same strain. It is well known that *S. cerevisiae* accumulates these carbohydrates at growth rates below 0.1 h^-1^ whereby the contents are related to the duration of the G1 phase [40]. Under carbon limited conditions trehalose and glycogen serve as carbon and energy reserves to enable the survival during starvation but are also mobilized to facilitate a transient increase in the ATP flux for progression through the cell cycle [41]. For all three strains the accumulations of trehalose and glycogen were strongly dependent on the cultivation temperature, with highest glycogen accumulation at 12 °C and highest trehalose accumulation at 39 °C. Increased trehalose accumulation at high cultivation temperatures have been observed before and were caused by a stimulation of trehalose synthase and inhibition of trehalose [42]. Because of the α-1,1-glycosidic bond linkage within the structure, trehalose has stronger resistance to heat and acid and was shown to be a preferable energy reserve over glycogen during stress conditions [37, 43], although the synthesis of trehalose requires more ATP per glucose than that of glycogen [36]. Besides, the degradation of trehalose releases two glucoses, whereas one glucose is released after the degradation of the α-1,4-glycosidic bond within glycogen. Apart from its role as a reserve carbohydrate, trehalose also has a protective function during stress conditions e.g. thermal stress, whereby it acts as the protector of membranes and proteins [44–46]. Therefore, the significantly higher trehalose accumulation of ADY5 and Ethanol Red might have contributed to the better growth performance at 39 °C compared with CEN.PK113-7D.

The superior growth performance of ADY5 during SBR cultivation at 12 °C conincided with a high capacity to accumulate glycogen. Increased carbohydrate accumulation, in particular glycogen, as a response to prolonged exposure of yeast to cold (10 °C) has been observed before [47]. Furthermore, a positive correlation between cell wall bound glycogen and viability under glucose deprived conditions was reported [48]. Although the precise function of this glycogen pool remains unclear, it might play a role in membrane stabilization which may improve the resistance to cold [49].

## 4. Conclusions

From a growth phenotypic screening of 12 industrial *Saccharomyces* strains for their temperature tolerance, we selected ADY5, Ethanol Red, and CEN.PK113-7D to further elucidate the possible underlying mechanisms for temperature tolerance. The chemostat results revealed significant differences in the metabolic response and cellular energetics between strains and among different growth temperatures. Despite of a fixed growth rate, different growth temperatures resulted in large differences between the three strains in terms of net conversion rates, substrate yields and energetic efficiency of biomass formation. All strains showed a decrease of protein content at supra-optimal temperatures which was nevertheless accompanied with a decrease of Y_ATP_, thus implying an increase of non-growth associated energy demands. Increased temperature tolerance coincided with higher energetic efficiency of cell growth, indicating that temperature intolerance is a result of energy wasting processes, such as increased turnover of cellular components due to temperature induced damage, e.g. protein misfolding. Further research is required to deepen our comprehension on the underlying mechanisms. With this knowledge, we can develop and apply strategies to obtain tailored cryo- and-thermo tolerant yeasts for industrial applications.

## 5. Methods and materials

### 5.1. Yeast strains, growth conditions, and storage

A total of 13 *Saccharomyces* strains used in this study of which one *S. uvarum*, one *S. cerevisiae* x *S. cerevisiae* hybrid and the others were *S. cerevisiae* species (Table 8). Inocula were prepared by introducing a single colony of a pure culture of each strain into 5 mL sterilized synthetic medium [50] with 15 g·L^-1^ C_6_H_12_O_6_·H_2_O in a 30°C incubator shaker at 220 *rpm*. Biomass stocks were prepared by the addition of sterilized glycerol to the exponentially growing cultures of all 13 strains, resulting the final concentration of 30% (v/v). The biomass stocks were store aseptically at −80°C. These frozen stocks were used to inoculate the different experiments described as below.

The growth profile can be obtained by growing the yeast strains on a microtiter plate at different temperatures ranging from 12°C to 40°C under aerobic condition. Pre-culture was grown at 30°C and 220 *rpm* in sterilized synthetic medium [50] with 7.5 g·L^-1^ C_6_H_12_O_6_·H_2_O. the pre-culture was transferred to the microtiter plate (24 wells) with fresh synthetic medium resulting an initial optical density at 600nm of approximately 0.1 and grown at the temperature ranged from 12°C to 40°C with continuous shaking (300 *rpm*, 1-inch amplitude). Growth was monitored via the optical density at 600 nm in a Synergy HTX Multi-Mode reader (BioTek, USA), and measurement was taken every 15 minutes for 18 hours. For the cultivation at temperature below 30°C, microtiter plates were cultivated outside the Synergy HTX Multi-Mode reader and measured the OD_600_ by the Synergy HTX Multi-Mode reader every 8 hours for 4 days. The growth profile of each strain at different temperatures was obtained by triplicate measurements and was fit to two models.

### 5.2. Primary model

The maximum specific growth rates of each strain at different temperatures were obtained by fitting the experimental OD_600_ to the corrected modified Gompertz model equation modified from the original version [25].

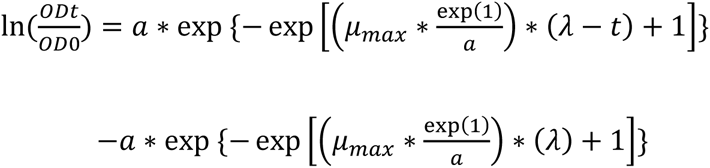

Where OD_0_ is the initial OD_600_ and OD_t_ is that at time t; *a* is the asymptotic maximum of 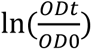; µ_max_ is the maximum specific growth rate with a unit of h^-1^, and λ is the lag phase period. All the parameters of time have a unit of hour.

### 5.3. Secondary model

The CTMI model was used to fit with the obtained µ_max_ of each strain at different temperatures. The CTMI has the following expression;

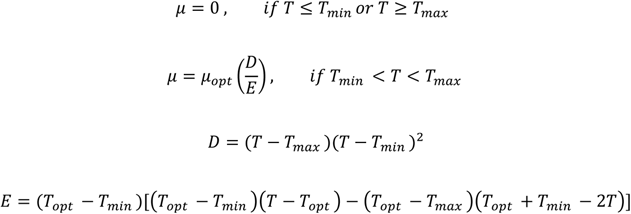

Where T_max_ is the temperature above which no growth occurs, T_min_ is the temperature below which no growth is observed, and T_opt_ is the temperature at which μ_max_ is equal to μ_opt_. Both the primary and secondary models were fitted by minimizing the residual sum of squares (RRS) with respect to the experimental data.

### 5.4. Fermentation set-up

All pre-cultures were grown aerobically at 220 rpm and at 30°C in the sterilized medium containing 5 g·L^-1^ (NH_4_)_2_SO_4_, 3 g·L^-1^ KH_2_PO_4_, 0.5 g·L^-1^ MgSO_4_·7H_2_O, 22 g·L^-1^ C_6_H_12_O_6_·H_2_O, 1.0 mL·L^-1^ of trace element solution, and 1.0 mL·L^-1^ vitamin solution [50]. The sterilization of the medium was performed using a 0.2 µm Sartopore 2 filter unit (Sartorius Stedim, Goettingen, Germany).

The reactor vessels were equipped with norprene tubing, to minimize the diffusion of oxygen into the vessels and were sterilized by autoclaving at 121°C. The exhaust gas from all fermentations was passed through a condenser kept at 4.0°C and then through a Perma Pure Dryer (Inacom Instruments, Overberg, The Netherlands) to remove all water vapour and subsequently entered a Rosemount NGA 2000 gas analyser (Minnesota, USA) for measurement of the CO_2_ concentration. The medium of all fermentations was continuously sparged with nitrogen gas prior and was contained 5.0 g·L^-1^ (NH_4_)_2_SO_4_, 3.0 g·L^-1^ KH_2_PO_4_, 0.5 g·L^-1^ MgSO_4_·7H_2_O, 22.0 g·L^-1^ C_6_H_12_O_6_·H_2_O, 0.4 g·L^-1^ Tween80, 10 mg·L^-1^ ergosterol, 0.26 g·L^-1^ antifoam C (Sigma-Aldrich, Missouri, USA), 1.0 mL·L^-1^ trace element solution, and 1.0 mL·L^-1^ vitamin solution [50]. The cultivations were carried out at temperatures of either 12.0 ± 0.1 °C, 30.0 ± 0.1°C or 39.0 ± 0.1 °C, by pumping cooled or heated water through the stainless-steel jacket surrounding the bottom part of the reactor vessel using a cryothermostat (Lauda RE630, Lauda-Königshofen, Germany). The water temperature of the cryothermostat was controlled by using the signal of a Pt 100 temperature sensor inside the reactor, for accurate measurement and control of the cultivation temperature. Anaerobic conditions were maintained by continuously gassing of the reactor with nitrogen gas at a flowrate of 1.00 ± 0.01 SLM (standard liter per minute) using a mass flow controller (Brooks, Hatfield, USA). Also, the feed medium was kept anaerobically by sparging with nitrogen gas. The nitrogen gas was sterilized by passing through hydrophobic plate filters with a pore size of 0.2 μm (Millex, Millipore, Billerica, USA). The culture broth in the reactor was mixed using one 6-bladed Rushton turbine (diameter 80 mm) operated at a rotation speed of 450 rpm. The pH was controlled at 5.00 ± 0.05 by automatic titration with 2.0 M KOH.

### 5.5. Sequential batch cultivations

The progression of all batch fermentations was monitored by the CO_2_ measurement in the exhaust gas from the off-gas analyser and the base addition into the culture broth. The base bottle was placed on a load cell (Mettler Toledo, Tiel, The Netherlands), and thus the amount of base addition was measured by the weight decreased from the base bottle and by a totalizer of a feed pump. When there was no based addition for a defined amount of time, the culture broth was automatically drained from the bottom of the bioreactor by the effluent pump until there was 0.20 kg left which was regulated by using the weight balance of the bioreactor (Mettler Toledo, Tiel, The Netherlands). Fresh medium was subsequently added to fill the reactor until the total volume was 4.0 kg. The volume of the broth in the bioreactor was regulated by the weight balance of the bioreactor (Mettler Toledo, Tiel, The Netherlands) which was placed underneath the bioreactor. Six sequential batches were carried out in the above-mentioned cycle at each temperature set point (12°C, 30°C, and 39°C). The temperature set point was changed after every 6 sequential batches. The maximum specific growth rate of each strain at each cultivation temperature was calculated and averaged from the CO_2_ off-gas profiles of the last three sequential batches.

### 5.6. Chemostat Fermentation

All chemostat cultivations were carried out at a dilution rate of 0.030 ± 0.002 h^-1^ in 7 L bioreactors (Applikon, Delft, The Netherlands) equipped with a DCU3 control system and MFCS data acquisition and control software (Sartorius Stedim Biotech, Goettingen, Germany).

All chemostat fermentations were initially operated in anaerobic batch cultivation with 4 L biomass culture broth in a 7 L bioreactor (Applikon, Delft, The Netherlands) to achieve enough biomass at the start of the chemostat phase. Four hundreds mL of pre-culture was inoculated in each batch cultivation. When the off-gas CO_2_ level from the batch cultivation dropped significantly close to the level after the pre-culture inoculation, this indicated the end of the batch phase. The fermentation was switched to a chemostat phase by switching on the continuous feed of the sterilized medium to the bioreactor, of which the sterile feed medium was pumped into the reactor vessel at a constant flowrate using a peristaltic pump (Masterflex, Barrington, USA), such that the outflow rate of the culture broth was 120 ± 1 g·h^-1^. The effluent vessel was placed on a load cell of which the signal was continuously logged for accurate determination of the dilution rate of the chemostat and manual adjustment of the medium feed rate if needed. The working volume was kept constant at 4.00 ± 0.05 kg using the weight balance of the bioreactor (Mettler Toledo, Tiel, The Netherlands) which controlled the effluent pump. All chemostat cultures reached a steady-state after 5 volume changes, which was apparent from stable CO_2_ levels in the exhaust gas and the biomass dry weight concentration were obtained. After reaching steady-state, triplicate samples at four sampling time points were withdrawn during another period of 4 to 5 volume changes, for quantification of the concentrations of biomass, residual glucose and extracellular metabolites.

### 5.7. Analytical methods

Optical density was monitored using a Libra Su spectrophotometer (Biochrom Libra, Cambridge, UK) at a wavelength of 600 nm. Biomass dry weight was determined using the filtration of sample broth over a dry nitrocellulose filter (0.45 μm pore size, Gelman laboratory, Ann Arbor, USA) which were dried in a 70°C oven overnight. After the filtration of the sample broth, two sample volume of demineralized water was used to wash the filters which were subsequently dried in the oven at 70°C for two days. Prior and after sample filtration the filters were measured after cooling down in a desiccator for two hours. Extracellular metabolite of the sample broth was obtained using cold stainless-steel beads [51]. The resulting supernatant broth was immediately frozen by liquid nitrogen and followed by the storage at −80°C. The supernatant broth was defrosted and analysed in duplicate using high-performance liquid chromatography (HPLC) with a Bio-Rad Aminex column (Bio-Rad Laboratories, California, USA) at 60°C. The column was eluted with 5.0 mM phosphoric acid at a flow rate of 0.6 mL·min^-1^. Ethanol and glycerol were detected with a Waters 2414 refractive index detector (Waters Corporation, Massachusetts, USA), while a Waters 1489 UV-Vis detector (Waters Corporation, Massachusetts, USA) was used to detect acetate, lactate, malate, and succinate. Residual glucose was measured by ion chromatography using Dionex-ICS 5000+ (Thermo Fisher Scientific, Massachusettts, USA).

### 5.8. Metabolic Flux Analysis and Data Reconciliation

The metabolic flux distributions as well as the best estimates of the biomass specific net conversion rates of the chemostat experiments were obtained via metabolic flux analysis using a stoichiometric model for anaerobic growth of *S. cerevisiae* [19]. With sufficient available conversion rates as input variables an overdetermined system was obtained, allowing to find the best estimates of the biomass specific net conversion rates within their error margins as well as the metabolic flux distributions under the constraint that the elemental and compound balances were satisfied [52, 53].

### 5.9. Total organic carbon and total nitrogen measurement

The total organic carbon (TOC) of the chemostat total broth and the chemostat supernatant broth was calculated from the subtraction between the total carbon (TC) and the total inorganic carbon (TIC) which were both measured by a total organic carbon analyser (TOC-L CSH, Shimadzu, Kyoto, Japan).

The total nitrogen (TN) of the freeze-dried biomass (10 mL culture broth) from the chemostat culture was measured by a total nitrogen unit (TNM-L, Shimadzu, Kyoto, Japan). The TN content of a sample were in the form of ammonium, nitrite, nitrate, as well as organic compounds.

The injection of the samples for both TOC and TN measurement was carried out by an auto-sampler (ASI-L, Shimadzu, Kyoto, Japan).

### 5.10. Cellular protein measurement

Thirty millilitres chemostat cultural broth was drawn from the reactor and subsequently centrifuged at 4°C and at 5000 *rpm* for 5 minutes. The supernatant was discarded, and the biomass pellet was immediately frozen in liquid nitrogen and stored in −80°C prior freeze-drying. The cellular protein of the freeze-dried biomass was determined using Biuret method as described in [54] at which freeze-dried BSA was used as standard.

### 5.11. Cellular glycogen measurement

Approximately 2 mg of biomass was quenched into 100% methanol, which was chilled in −40°C prior, using a rapid sampling setup [55]. The quenching volume ratio of sample broth and methanol was 1 to 6. The quenching sample in 100% methanol was centrifuged at −19°C and at 5000 *rpm* for 5 minutes. The supernatant was discarded, and the biomass pellet was immediately frozen in liquid nitrogen and stored in −80°C.

The biomass pellet was washed twice with 1.5 mL cold Mi-Q water in an Eppendorf tube and followed by centrifugation at 4°C and at 8000 *rpm* for 2 minutes. The supernatant was discarded. The pellet was dissolved in 250µL of 0.25 M sodium carbonate solution and subsequently incubated at 95°C for 3 hours with continuously shaking. After the incubation, 600 µL of 0.2M sodium acetate was added into the mixture, the pH of which was adjusted to 5.3 with 1M acetate acid afterwards. The hydrolysis of the glycogen was performed by adding α-amyloglucosidase dissolved in 0.2 M sodium acetate to the mixture to have the final concentration of 1.2 U/mL. The reaction of the hydrolysis was carried out at 57°C overnight with continuously shaking. The equivalent glucose released from the glycogen digestion was determined in triplicate using the UV bioanalysis kit (R-Biopharm/Roche, Darmstadt, Germany). The resulting assay was measured at the wavelength 340 nm by a spectrophotometer.

### 5.12. Cellular trehalose measurement

Approximately 2 mg of biomass was quenched into 100% methanol, which was chilled at −40°C prior, using a rapid sampling setup [55]. Sample filtrate was washed twice with 20 mL 80% (v/v) methanol, which was chilled at −40°C prior, and was extracted with boiling ethanol as described in [55]. Cellular trehalose of the chemostat culture was measured by GC-MS analysis as described in [56]. C^13^ labelled cell extract was added into the extracted sample as internal standard [57].

## Acknowledgements

We would like to thank Judith Cohen and Kristen H. David for technical supports in the chemostat fermentations. This research is acknowledged to the project ERA-IB “YeastTempTation” (ERA-IB-2-6/0001/2014) and to the Portuguese Foundation for Science and Technology (FCT) under the scope of the strategic funding of UID/BIO/04469/2020 unit and to BioTecNorte operation (NORTE-01-0145-FEDER-000004) funded by the European Regional Development Fund under the scope of Norte 2020 – Programa Operacional Regional do Norte. Lallemand Ibéria, SA supplied the industrial yeast strains in this research.

## Author Contributions

All authors have been involved in the design of the experiments. CEC,EGR, and KYFL performed the growth phenotypic screenings in microtiter plates. EGR and KYFL performed the computational simulations. EGR performed the HCL analysis. KYFL performed the measurements of total proteins, total nitrogen, and storage carbohydrates of the chemostat cultures. KYFL and WVG performed the fermentations in bioreactors, analysis of the results, and wrote the draft of the paper. CEC and LD made the graphic abstract. All authors read, edited, and approved the final manuscript.

## Supplementary materials

**Table S1.** Average acetic acid concentrations (mmol·L^-1^) in the extracellular broth during steady-state of chemostat cultivation of the three strains at 12 °C, 30 °C, and 39 °C. Standard errors were obtained from four measurements at different time points during the steady-states.

|  | 12°C | 30°C | 39°C |
| --- | --- | --- | --- |
| ADY5 | 0.00 ± 0.00 | 0.00 ± 0.00 | 1.06 ± 0.10 |
| CEN.PK113-7D | 2.84 ± 0.10 | 0.52 ± 0.04 | 0.54 ± 0.09 |
| Ethanol Red | 1.16 ± 0.03 | 0.67 ± 0.12 | 1.19 ± 0.03 |

**Table S2.** Average cellular nitrogen contents (g total N·g_DW_^-1^) of the three strains during anaerobic steady state chemostat cultivation at 12 °C, 30 °C, and 39 °C. Standard errors were obtained from four measurements at different time points during the steady-states.

|  | 12°C | 30°C | 39°C |
| --- | --- | --- | --- |
| ADY5 | 0.060 ± 0.004 | 0.046 ± 0.004 | 0.044 ± 0.002 |
| CEN.PK113-7D | 0.054 ± 0.003 | 0.051 ± 0.002 | 0.044 ± 0.003 |
| Ethanol Red | 0.061 ± 0.005 | 0.067 ± 0.001 | 0.046 ± 0.001 |

**Table S3.** First order evaporation constants for broth (water + ethanol) and ethanol determined from measured broth and ethanol disappearance of an ethanol water mixture during fermentation conditions at 12 °C, 30 °C, and 39 °C.

| Temperatures | $k_{\text{broth}}$<br>$\frac{g_{\text{broth evaporated}}}{(kg_{\text{broth in bioreactor}} \cdot h)}$ | $k_{\text{eth}}$<br>$\frac{mmol_{\text{ethanol evaporated}}}{(mol_{\text{ethanol in broth in bioreactor}} \cdot h)}$ | p-value |
| --- | --- | --- | --- |
| °C |  |  |  |
| 12 | -0.075 ± 0.009 | -1.62 ± 0.20 | 0.260 |
| 30 | -0.332 ± 0.010 | -4.44 ± 0.31 | 0.228 |
| 39 | -0.359 ± 0.013 | -7.71 ± 0.22 | 0.293 |

